# A novel setup for simultaneous two-photon functional imaging and precise spectral and spatial visual stimulation in *Drosophila*

**DOI:** 10.1101/2020.03.26.009688

**Authors:** R. C. Feord, T. J. Wardill

**Affiliations:** Physiology, Development & Neuroscience, University of Cambridge, CB2 3EG, UK; Ecology, Evolution and Behavior, University of Minnesota, St Paul, MN, 55108, USA

## Abstract

Motion vision has been extensively characterised in *Drosophila melanogaster*, but substantially less is known about how flies process colour, or how spectral information affects other visual modalities. To accurately dissect the components of the early visual system responsible for processing colour, we developed a versatile visual stimulation setup to probe combined spatial, temporal and spectral response properties. Using flies expressing neural activity indicators, we tracked visual responses in the medulla to a projected colour stimulus. The introduction of custom bandpass optical filters enables simultaneous two-photon imaging and visual stimulation over a large range of wavelengths without compromising the temporal stimulation rate. With monochromator-produced light, any spectral bandwidth and centre wavelength from 390 to 730 nm can be selected to produce a narrow spectral hue. A specialised screen material scatters each band of light across the visible spectrum equally at all locations of the screen, thus enabling presentation of spatially structured stimuli. We show layer-specific shifts of spectral response properties in the medulla correlating with projection regions of photoreceptor terminals.

## INTRODUCTION

The fruit fly, *Drosophila melanogaster*, is a key model for invertebrate vision research ^[1]^. The small diameters of cells in the early visual neuropil have long limited electrophysiological approaches to recording neural activity in the optic lobes, a problem recently overcome with advances in technology and genetic tools that make functional imaging methods possible ^[2–4]^. Though these imaging methods have facilitated extensive characterisation of motion vision in flies ^[5]^, how flies process colour information, or how the spectral content of light affects other visual modalities, remains largely unknown.

Photoreceptors in the *Drosophila* eye express different classes of photoreceptive cells, referred to as R1-R8. Within an ommatidium, the six outer photoreceptors (R1-R6) express the broadband rhodopsin (Rh) 1. The more centrally located cells (inner photoreceptors), R7 and R8, express Rh3-6 photopigments, which are narrowly tuned to UV, blue and green bands of the spectrum^[6–9]^. Additional parameters, notably screening pigments, exert influence on the spectral response properties of the photoreceptor cells^[10]^. Light-absorbing screening pigment encircling each ommatidium restricts off-axis light from reaching the photoreceptors ^[10]^ and yellow-coloured pigment granules in R1-R6 cells contribute to spectral tuning via a dynamic and light-dependent migratory pattern ^[11, 12]^.

The overall picture of colour vison in *Drosophila* is being pieced together with improved knowledge of photoreceptors, candidate cells involved in circuitry, and colour-guided behaviour ^[13–19]^, but much still remains to be characterised. The long-standing dogma advocates that the motion and colour vision processing streams are neatly separated, with R1-R6 providing input to the motion detection pathway and R7-R8 providing input to the colour vision pathway ^[13, 20–22]^. More recent evidence, however, demonstrates that signals arising from R7 and R8 improve motion processing in *Drosophila* ^[23]^. This paradigm of combining modalities at an early stage of visual processing has previously been demonstrated as a strategic mechanism for improved perceptual discrimination and retention of salient visual features ^[24–26]^.

Commercially available visual displays are tailored to the primate retina. Their three-colour channel configuration provides an excellent experience to the trichromatic user but fall short for visual systems with different photoreceptor sensitivities. Furthermore, the high flicker-fusion frequencies in invertebrates, beyond 120 Hz in *Drosophila melanogaster* ^[27]^, outpace the refresh rates of commercial displays. An array of experimental systems for visual stimulation have been developed, but many constraints still limit the investigation of combined colour and motion processing. The production of bands of monochromatic light is commonly achieved via a broadband light source coupled either to a monochromator ^[8]^, individual colour filters ^[28, 29]^, LEDs ^[30]^, or more recently LED-based monochromator systems ^[31]^. Such monochromatic light provides full-field stimulation but lacks any spatial structure. Paradigms for spatially patterned stimuli include LCD displays, LED panels or projectors, none of which offer the option of many colours ^[17, 32–34]^. In order to probe visual response properties to combined modalities, the integration of spectral and spatial resolution within a stimulation paradigm is essential. In addition, the spectrally broad and high detection sensitivity of two-photon imaging systems restricts the visual stimuli’s spectral range and intensity: light applied within the detection range of the microscope will result in an artefact on the acquired image consequently restricting the range of wavelengths available for visual stimulation.

In order to determine the precise contribution of spectral information to visual computations, whether general or colour-specific, we designed a system that offers fine-wavelength resolution across a large portion of the spectrum, that can be calibrated to produce isoluminant stimuli over a biologically-relevant range of intensities and that allows the presentation of spatially- and temporally-structured stimuli. This system presents several advantages not offered in any previous imaging-based setups for characterising physiological responses to colour. These include (1) the delivery of coloured stimuli simultaneous to the acquisition of two-photon images (previous methods rely on the scanner fly-back periods for stimulation ^[17, 34]^, (2) the ability to arbitrarily chose the centre wavelength and bandwidth), and (3) a rear projection screen which maintains isoluminant hues across the visible spectrum. Using this setup, we characterised intensity-response relationships and spectral response profiles of the pan-neuronally labelled medulla in several genetically modified strains of *Drosophila*, varying in opsin functionality and screening pigment density.

## RESULTS

### A modified monochromator-projector-microscope system

A visual stimulus was presented to a fly while imaging its neural responses via a two-photon microscope. To produce a range of colours (selectable narrow bands of the spectrum), we modified a projector to use a monochromator as its light source (**Figure 1A**). We introduced custom Semrock bandpass optical filters to the imaging and visual stimulation pathways (**Figure 1B**). These filters allowed for the presentation of a visual stimulus with minimal contamination to the calcium signal read. One filter set, added to the monochromator, blocks the wavelengths of light within the microscope’s detection range. The other set of optical filter combination is integrated into the microscope to limit the wavelengths of light entering the detectors and works to reject bleed-through of detectable light wavelengths from the visual stimulus by means of an arrangement of high optical density bandpass filters (further details of the filters in **Supplementary Figure S1**).

**Figure 1.**
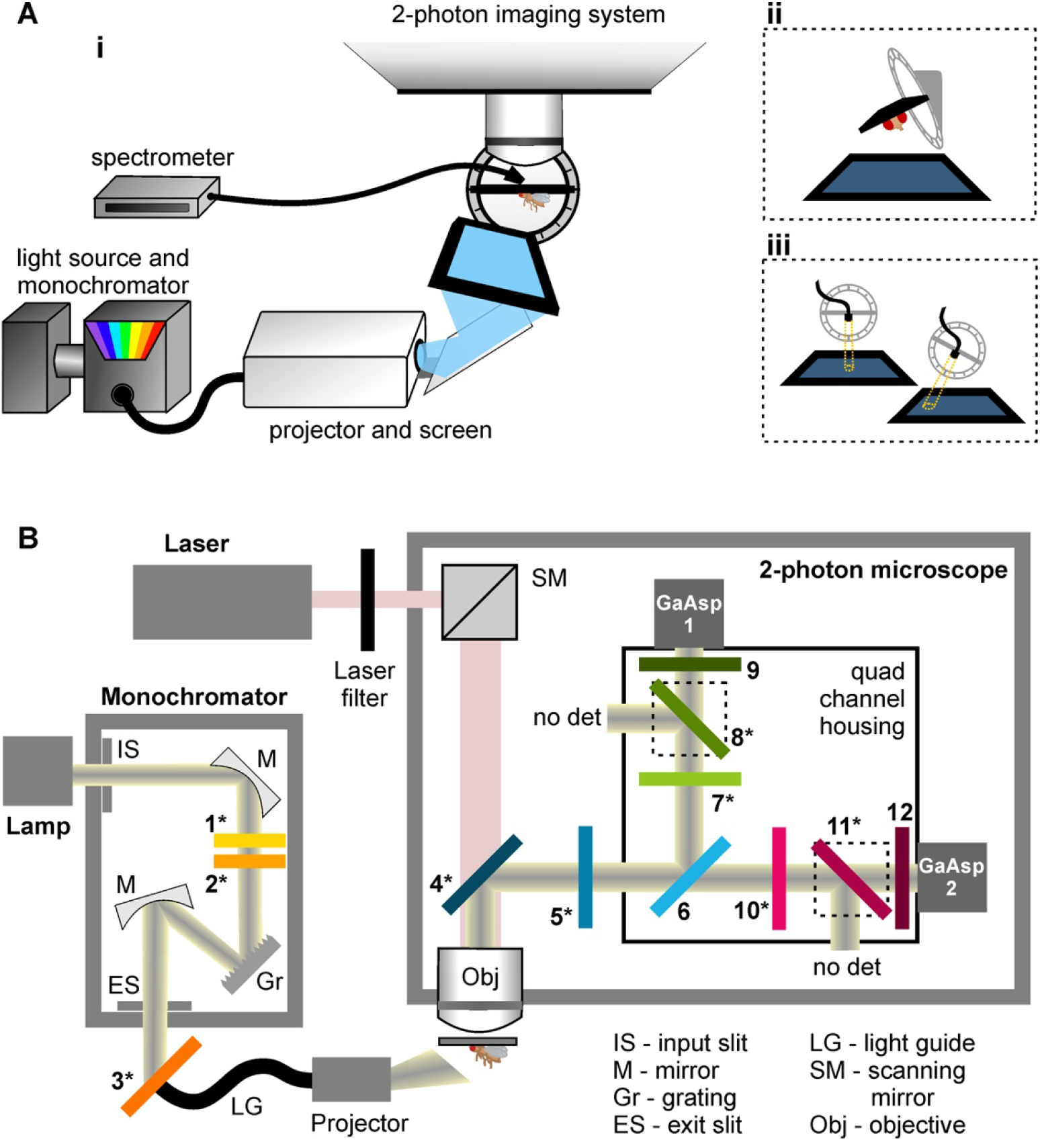
A novel setup enabling simultaneous two-photon functional imaging and precise colour visual stimulation. (**A**) Placement of the fly in the holder with its cuticle removed to expose the optic lobes allows simultaneous neural activity imaging and visual stimulation. (**Ai**) A modified projector with a monochromator light source projects over 50 different bands of the visual spectrum onto a screen. These colour bands are precisely calibrated using a spectrometer to measure the radiance value of the visual stimulus. Both the monochromator and the spectrometer are jointly controlled: automated calibration of the stimulus produces the required light intensity and spectral content via a feedback loop. A tilted holder allows (**Aii**) the spectrometer position to be adjusted for measurements of specific points of the screen, as well as (**Aiii**) to maximise the coverage of the fly’s eye by the screen. (**B**) Modifications of the optical pathway in the monochromator and the microscope are necessary to increase the bandwidths of stimulus wavelengths without detection by the microscope. Light from a broadband tungsten 150W bulb (lamp) is selectively transmitted through the monochromator via an input slit (IS), several mirrors (M), a grating (Gr) and an exit slit (ES). Three custom Semrock filters are added along this pathway. Filters **1** and **2** are long-pass filters that prevent the transmission of harmonics in the UV range. Filter **1** is moveable and is only used for light above 460 nm. Filter **3** prevents the transmission of the bands of red and green light that are detected by the microscope. A custom Semrock filter (**4**) replaces the dichroic mirror at the start of the imaging pathway, and combined with **5** and **6**, these three filters serve to block the transmission of any light beyond the narrow bands of red and green detected by the microscope. A further three filters are combined for each GaAsP detector. Filters **7** and **10** block light that falls outside the red and green band ranges. Filters **8** and **11** are long-pass filters. Finally, filters **9** and **12** selectively transmit only specific red and green band of light for GaAsP detectors 1 and 2 respectively. Unlike classic quad housing designs, there are only two detectors instead of four and 200-500 nm and 550-600 nm light along the GaAsp1 and GaAsp2 detector pathways respectively is discarded. Filters denoted with asterisks (*) are a custom addition to the commercially available version of the system. Filter spectra are reported in **Supplementary Figure S1**.

### Combined spectral and spatial precision

The light produced by the monochromator can be modified for two parameters, wavelength and intensity, providing ample versatility to produce stimuli across the visible spectrum and across several log units of intensity. To test for spectral response independent of brightness, all monochromatic bands of light were calibrated to produce the same radiance with minimal variation over time (**Supplementary Figure S2**). Bands spanned from 385 to 725 nm at approximately 5 nm centre wavelength intervals. Gaps in the transmitted light between 500-540 nm and 610-650 nm (**Supplementary Figure S2**) exist because of the dichroic filters integrated to the monochromator. Furthermore, we sought to ensure minimal variation of projected light across the two-dimensional plane. As the distance between the fly’s eye and the screen increases towards the outer edges of the screen, the brightness diminishes accordingly (**Figure 2A**). A more problematic source of variation, however, stems from light scattering. Scattering is an inherent and necessary property of the screen to ensure light diffuses over the array of angles required to reach the fly’s eye (**Figure 2A**). Scattering properties of a material, however, are coupled to the wavelength of the light. This creates the challenge of identifying a screen material that scatters light in an equivalent manner across the UV and visible spectrum. We identified a screen fitting these requirements (Da-Lite, Polacoat^®^ Flex Plex Video Vision), ensuring that calibrated isoluminant bands of light at the screen centre retain their flat isoluminant calibration across the screen from 385 to 720 nm (**Figure 2B** and **2C**). A small variation between the UV and red light occurs at the outer edges of the screen. However, by delimiting a circular ellipse (diameter: 400 pixels/38.8 degrees of visual field), with its centre realigned to correspond to maximal brightness (blue dot, **Figure 2B**), the projected stimulus retains spectral constancy (**Figure 2D**).

**Figure 2.**
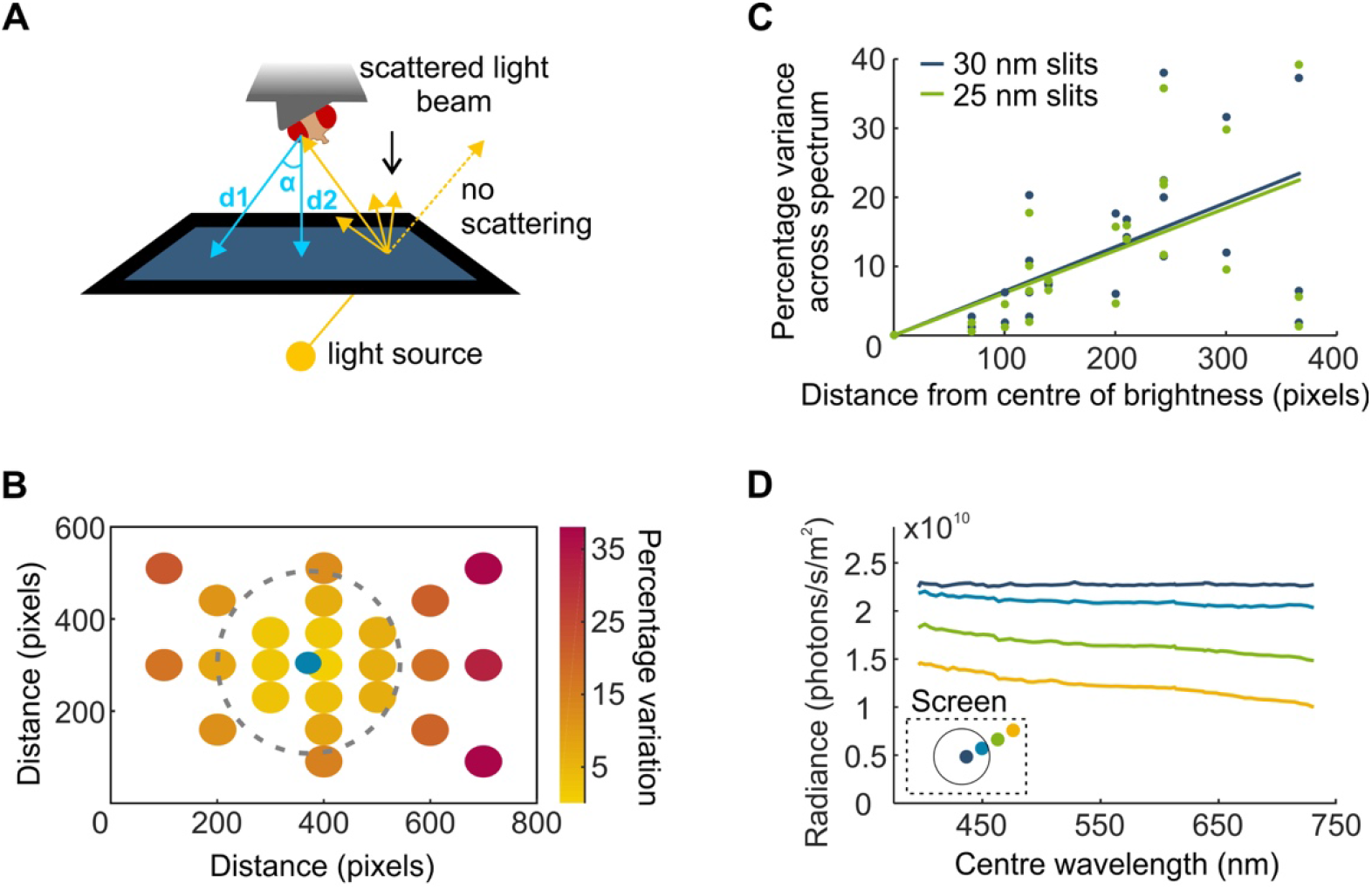
Optical properties of the screen allow for combined high spatial and high spectral precision. (**A**) Rear-projection of the stimulus from a point light source onto a screen causes variation in light intensity across the surface (brightest at the centre and decreased towards the edges). Light scattering is a necessary property of the screen material to ensure light reaches the fly’s eye despite its angle variance to the screen. However, scattering properties are wavelength-dependent perturbing spectral constancy across the screen. (**B**) The percentage variation between minimum and maximum radiance curve integrals from an isoluminant calibration of wavelength bands between 385 nm and 730 nm (385 nm light at screen centre approx. 2.35×10^10^). This variation is measured for 24 different screen locations along the orthogonal and diagonal axes of the screen. A 400-pixel diameter circle delineated by the dotted line corresponding to 40 degrees of visual field encompasses a minimally-varying portion of the screen. The blue dot represents a small shift of the circle centre relative to the screen centre as maximal brightness is not perfectly centred. (**C**) The percentage decrease in intensity from the screen centre plotted as a function of distance for a brighter (385 nm light at screen centre approx. 2.35×10^10^ and a dimmer (385 nm light at screen centre approx. 1.42×10^10^ calibration. A linear regression fitted to the data shows the two intensities exhibit similar trends for variance across the spectrum. (**D**) Example traces of a flat calibration at the screen centre, measured along one diagonal of the screen. Minimal variation occurs within the circle specified in (B).

### Proof of concept: spectral response properties in the *Drosophila* medulla

#### General medulla responses

To probe spectral response properties in the medulla of *Drosophila*, we established intensity-response relationships and spectral response profiles by means of calcium activity indicator imaging in pan-neuronally labelled flies. Transgenic fly strains differed from each other for one or more of the following parameters: screening pigment density (red: high, orange: low), photoreceptor function (intact or Rh1-only) and calcium activity indicator (GCaMP6f: green, RGECO: red, **Figure 3A**).

**Figure 3.**
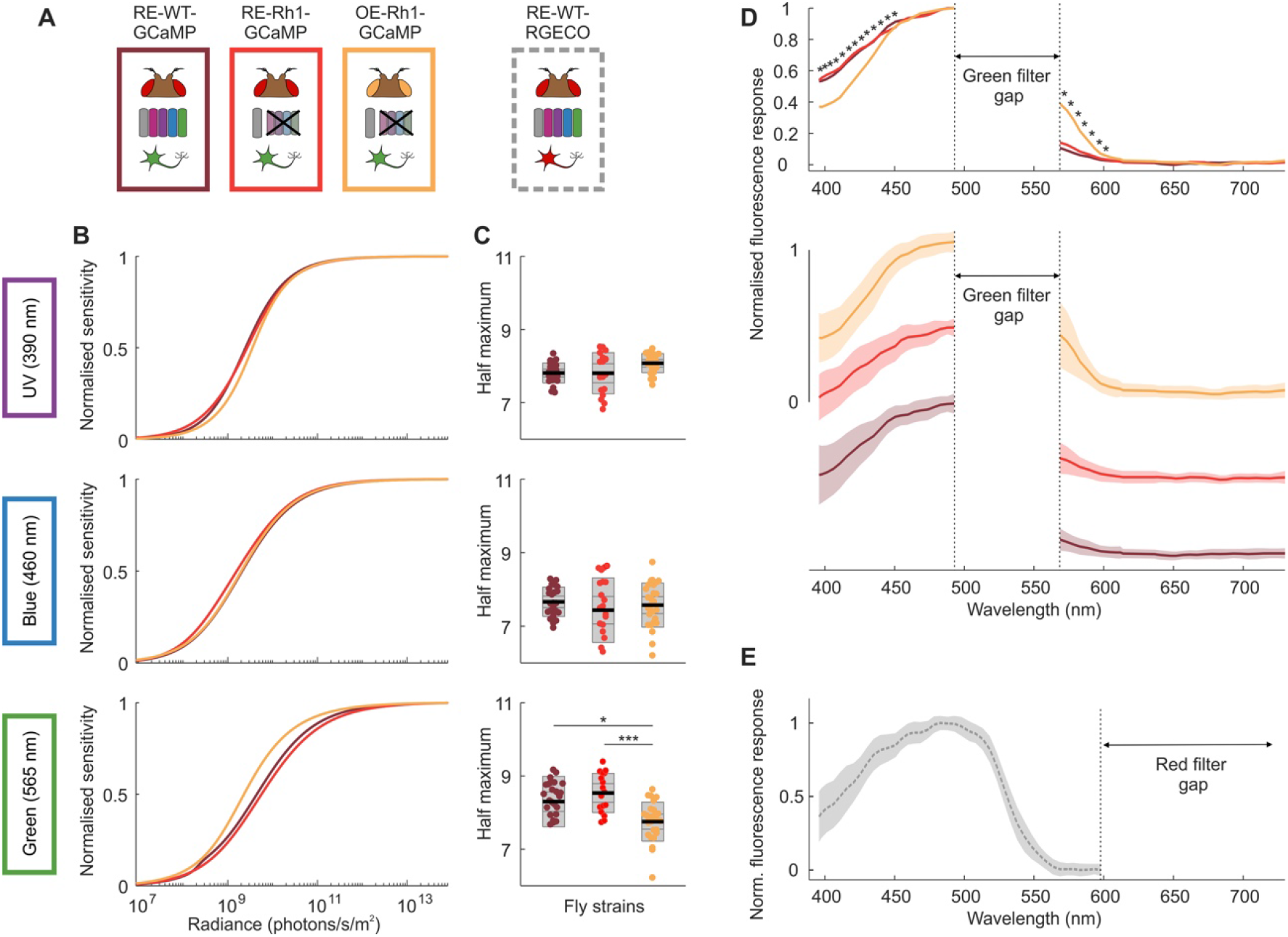
Reduced screening pigment expression biases the fly’s spectral sensitivity towards longer wavelengths of light. (**A**) Experiments were performed in four different fly strains expressing pan-neuronal calcium activity indicators – red eye, wildtype photoreceptors and GCaMP6f (RE-WT-GCaMP, bordeaux); red eye, Rh1 only and GCaMP6f (RE-Rh1-GCaMP, red); orange eye, Rh1 only and GCaMP6f (OE-Rh1-GCaMP, orange) and red eye, wildtype photoreceptors and RGECO (RE-WT-RGECO, grey). Intensity-response relationship curves (**B**) were established for three different bands of monochromatic light (UV, blue and green; centre wavelengths 390, 460 and 565 nm respectively) and the half maximum values (**C**) were extracted from fitted curves. Individual data points correspond to the average of ROI responses across an individually discernible layer structure (see methods) for a given fly preparation. Black line = mean, inner grey box = SEM, outer grey box = SD. Significant differences are noted with star values, with the P values from left to right: *P = 0.0105 and ***P = 0.0002 (one-way ANOVA) (**D**) Normalised spectral response profiles, determined by a sweep of equal intensity light pulses across the spectrum. Superimposed means-only traces are represented in the upper panel and the same mean traces are depicted staggered with the addition of standard deviations in the lower panel. Stars indicate statistically significant differences between fly strains for a given centre wavelength, further statistical information is reported in Figure S4 and supplementary table S1. For GCaMP-expressing flies, a gap in the spectral sweep is necessary in the green wavelengths of light corresponding to the microscope’s green detection range. (**E**) Mean trace ± SD of the spectral response profile for RGECO-expressing flies. The spectral sweep gap for RGECO-expressing flies sits at the red end of the spectrum.

A set of full-field light pulses of increasing intensity were applied to the fly’s eye (**Supplementary Figure S3**) and sigmoid curves fitted to establish intensity-response relationships for UV, blue and green light (390, 460 and 565 nm centre wavelength respectively, **Figure 3B**). We observed a left shift of the green intensity-response curve and half-maximum values in orange-eye flies by comparison with their red counterparts (**Figure 3B** and **3C**), indicative of increased sensitivity to longer wavelengths of light. Next, we applied a series of light pulses ranging from UV light through to red light set at approximately 5 nm centre wavelength intervals (**Supplementary Figure S3**) and plotted normalised spectral response curves (**Figure 3D**). Our results revealed strain-specific sensitivity to colours: orange-eye flies exhibited a decreased sensitivity in the blue range and an increased sensitivity in the green range by comparison with their red eye counterparts (**Figures 3D** and **Supplementary Figure S5**). Coefficient values extracted from fitted intensity response curves (slope and half maximum, **Supplementary Figure S4**) suggest minor response property discrepancies between flies expressing GCaMP6f and RGECO (**Supplementary Figure S4**), attributable to differences in amplitude and decay times ^[35]^. Consequently, we did not pool data from RGECO-expressing flies with GCaMP6f data for statistical analyses. Nonetheless, this red-emitting indicator serves the valuable purpose of completing the spectral profile as the red GaAsP is used to record RGECO signals, thus allowing the GCaMP-restricted green wavelengths of the spectral sweep to be filled in (**Figure 3E**).

#### Layer-specific responses

Pan neuronal labelling of the medulla reveals clearly discernible and identifiable layer structures (**Figure 4A**) allowing us to delve into layer-specific responses for the intensity-response relationships and spectral response curves as above (**Figures 4C** and **Supplementary Figure S6**). Our results demonstrate that half maximum values for UV, blue and green light exhibit layer-specific variability between fly strains (**Figure 4C**). Although most layer groupings retain the higher sensitivity of orange-eye flies to green light, this difference disappears in layers M2 and layers M6-M7. The decreased sensitivity of orange-eye/Rh1-rescue flies relative to the red eye/wildtype photoreceptor flies in the UV-blue range of the spectral response curve (**Figure 3D**) is only apparent in layers M6-M7 and M9-M10. Strikingly, the red eye/Rh1-rescue flies exhibit increased sensitivity to UV, blue and green light in M2, but this increased sensitivity does not occur in the orange eye/Rh1 rescue flies. We also see this trend in layer M3, and we attribute the absence of statistical significance to the lower number of ROIs with successfully fitted intensity-response curves as response size is smaller than other layers.

**Figure 4.**
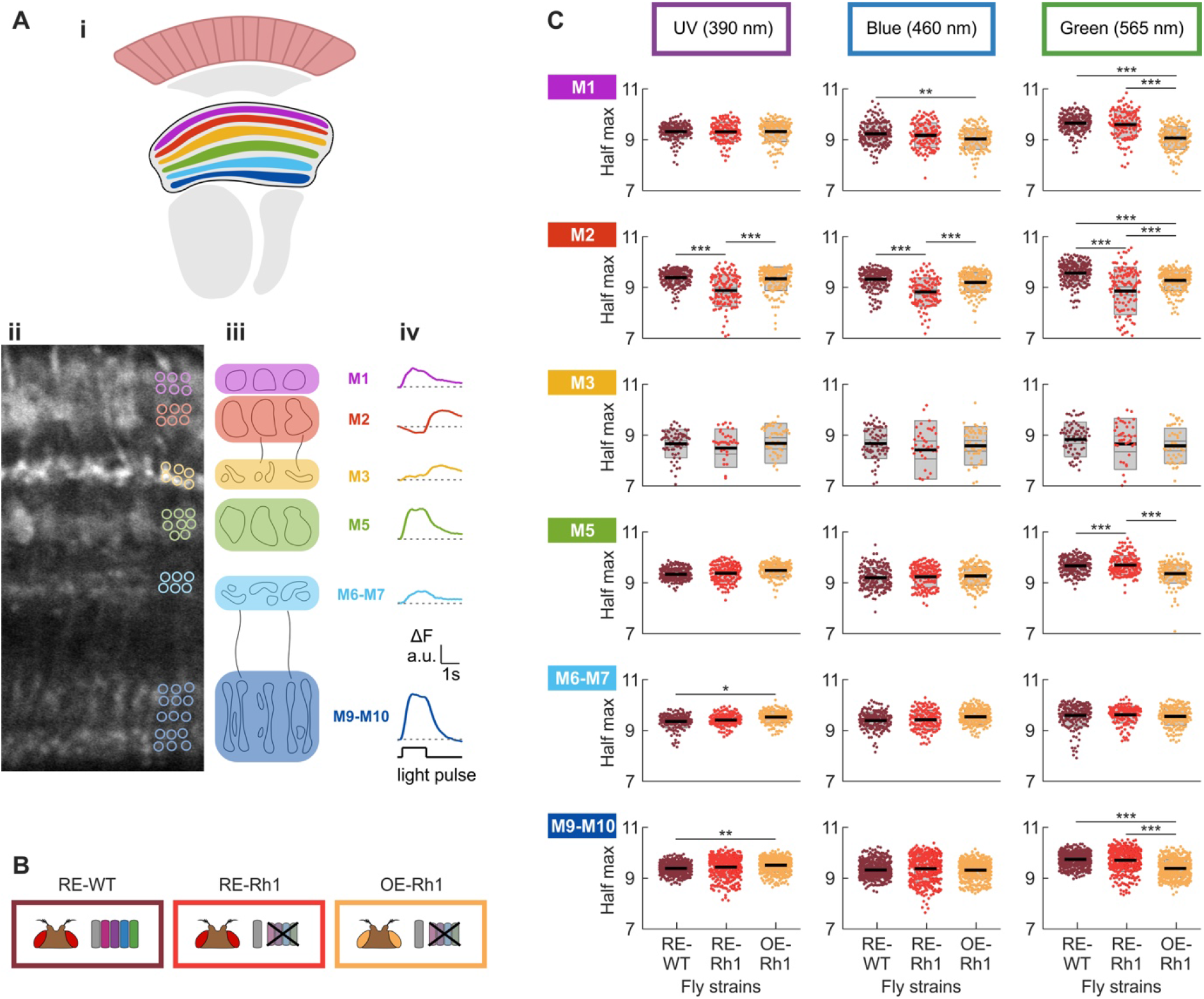
Spectral response properties vary across medulla layers. (**A**) Pan-neuronal GCaMP labelling of the medulla reveals six distinguishable layer structures (**Ai**, schematic; **Aii**, 2-photon image of baseline fluorescence), each exhibiting discernible cartridge-width substructures (**Aiii**). Response profiles of ROIs (segmentation examples shown in **Ai**) to a blue (440 nm) light pulse are stereotyped within a layer structure. These vary in temporal profile and size of response across layers M1, M2, M3, M5, M6-M7 and M9-M10 (**Aiv**). (**B**) Fly strains expressing pan-neuronal GCaMP6f– red eye/wild type photoreceptors (RE-WT, bordeaux); red eye/Rh1 only (RE-Rh1, red) and orange eye/Rh1 only (OE-Rh1, orange). (**C**) Half maximum values extracted from fitted intensity-response curves for three different bands of monochromatic light (UV, blue and green; centre wavelengths 390, 460 and 565 nm respectively) across six layer-groupings of the medulla as determined in (A). Black line = mean, inner grey box = SEM, outer grey box = SD. Significant differences are noted with star values, with one-way ANOVA P values reported in supplementary table S2.

## DISCUSSION

We developed a novel experimental setup that enables simultaneous two-photon functional brain imaging and visual stimulation across the spectrum in *Drosophila*. With this system, a fly can be presented with over 50 different wavelength bands of the visual spectrum, causing minimal disruption to the image quality produced by the microscope. Furthermore, our visual stimulation paradigm provides combined high spatial resolution and high spectral resolution, overcoming the customary trade-off between the two parameters.

The modifications to the optical pathway in both the monochromator and the microscope overcome the major limitation for spectrally-varied stimuli with two-photon imaging. The study of motion vision in *Drosophila* is usually carried out using blue light for visual stimulation, which offers an adequate wavelength range for Rh1-derived visual responses that sits outside of the PMT’s detection range. For the use of spectrally diverse stimuli, a workaround method was developed whereby the visual stimulus delivery occurs during the two-photon scanner fly-back period ^[17, 18]^. This approach, however, is limited by the discontinuous nature of the visual stimulus, which could introduce aliasing problems with moving patterns, but also reduces the microscope photon collection time thereby reducing the imaging sensitivity (this particularly affects calcium indicators with slow temporal responses). The introduction of custom Semrock bandpass filters in our system results in an extensive range of possible wavelengths for visual stimulation simultaneous to the acquisition of imaging data by the microscope. A gap in the spectrum corresponding to each PMT’s detection range still exists, but is considerably narrowed, and can be overcome by switching between neural activity indicators with different emission spectra detected by alternate PMT detectors.

In addition to the combination of simultaneous spectral visual stimulation and two-photon imaging, we also present a screen material that maintains near-constancy of the brightness of a wavelength band at all points of the screen from UV through to red light. To our knowledge this, surpasses all currently published screen materials in this regard, and offers a host of possibilities for characterisation of combined visual modalities such as motion and colour. The inevitable drop in radiance and spectral constancy towards the outer edges of the screen can be eliminated by restricting the stimulus to the central portion of the screen. Alternatively, if some variation of the brightness and spectral properties of the stimulus is not deemed to impact response properties, use of the extended screen area provides a greater coverage of the visual field.

Our understanding of the spectral quality of visual information transmitted by the different photoreceptive cells to downstream visual neurons is primarily based on the generation of spectral sensitivity curves for opsins or electroretinogram measurements of the surface of the eye ^[8, 36]^. A recent study, however, demonstrated that R7 and R8 terminals interact to transform the inner photoreceptor output to a biphasic sensitivity curve ^[17]^, exemplifying the modification of spectral tuning at the early stages of visual processing. Very little is known about how outputs of different photoreceptors might be combined by colour-encoding circuits, highlighting the need for high resolution spectral stimulus capabilities. Our system uses narrow bands of monochromatic light well suited to our investigation into spectral response properties, but it is entirely possible to modify the light source to produce multispectral light using a system such as the LED-based monochromator ^[31]^. Methods commonly employed in mammalian vision research, such as the sophisticated combination of multispectral light to effectively target only one class of photoreceptors ^[37]^, are limited in invertebrates. Further characterisation of the effect of pre-receptoral filtering, such as eye pigment screening, as well as instability and adaptation properties of visual pigments, is required. With this information, a smaller, but carefully selected, array of colour channels can be coupled to a display technology (screen or panel) such as the modified projector system with a five-primary light engine used in mouse vision research ^[33]^ or the arbitrary-spectrum spatial visual stimulator presenting up to six chromatic channels ^[34]^.

An ideal spectral stimulation range would extend further into the UV portion of the spectrum to match the known spectral sensitivities of Rh1, Rh3 and Rh5. Monochromators providing this range of UV light exist. However, the high-energy photons of the shorter wavelengths are lost by transmission throughout the optical pathway, with the DLP chip in the projector, in particular, drastically reducing the UV content of light. The ongoing progress in the development of UV transmitting projectors (e.g. Texas Instruments) and optics might soon provide an adequate and affordable solution and extend the capabilities of our system to include UV visual stimulation.

The experimental setup we describe here enabled us to record the responses to a range of narrow bands of light across the spectrum, over a range of intensities spanning several log units of light in the medulla. We find several modifications of spectral response properties in the summed activity of the pan-neuronally labelled neuropil between transgenic fly strains, demonstrating the precision, reliability and sensitivity of our setup. A reduction of screening pigment density results in increased sensitivity of medulla neurons to longer wavelength of light, consistent with the Goldsmith’s recruitment hypothesis and prior ERG recordings in flies ^[38–40]^.

Assessment of inter-layer variability revealed a diversity of responses. Most intriguing is the increased sensitivity of layers M2 and M3 in red eye flies that have lost R7 and R8 function, suggestive of inhibitory control of postsynaptic neurons in wild type flies by the R8 cells that terminate in M3 ^[41]^. Several reasons may explain why orange eye flies lacking functional photoreceptors do not also exhibit increased sensitivity in M2 and M3. Flies homozygous for the white null allele 1118 are known to have retina degeneration ^[42]^ and this may not be rescued with the mini-white vector expression. This effect could also be attributed to prolonged depolarisation afterpotential (PDA) ^[43]^, more common in white-eye flies ^[44]^, where the temporary light insensitivity of the photoreceptor requires red to shift metarhodopsin back to rhodopsin. This could also explain the decreased responsiveness to short wavelengths observed in the spectral response profile of this strain. The decreased sensitivity in the orange-eye flies to UV light in layers M6-M7 could be attributable to the lack of input from UV-sensitive R7 photoreceptors that project to this layer ^[41, 45]^. It is less clear, however, why these flies lose their increased sensitivity to green light uniquely in this layer structure. If the shift in response properties is caused by the photoreceptor terminals, green light should inhibit R7 activity ^[17, 18]^, resulting in decreased sensitivity in the red-eye flies expressing all functional opsins while maintaining unaffected high sensitivity in the orange-eye strain. Our data appears to suggest layer-specific shifts in spectral response properties in the medulla corresponding to projection regions of photoreceptor terminals. We did not detect summation of R7-R8 with R1-R6 signals proposed to drive lobular plate responses reported previously ^[23]^ as we found no difference between wild type and Rh1-only red-eye flies in most layers, however further investigation is required to establish this with certainty.

Our setup overcomes many limitations that have hindered the study of colour vision and the integration of visual modalities in *Drosophila*. Going forward, it will serve as a highly versatile tool to further recent breakthroughs in the field ^[17, 18]^ and unravel the neural circuitry underlying spectral processing.

## MATERIALS AND METHODS

### Fly stocks

Expression of the fluorescent calcium indicators GCaMP6f ^[3]^ and RGECO ^[35]^ was achieved using the GAL4/UAS expression systems ^[46]^. We used the pan-neuronal promoter nSyb (Bloomington Stock Center, 39171) and the following stocks for the red-eye norpA^36^ X chromosome mutant ^[23]^, white-eye norpA^36^ mutant (Bloomington Stock Center, 52276) and rhodopsin 1 rescue construct for norpA (Bloomington Stock Center, 9048). Four different fly stocks were used in this study:

1. w[+]; P{yw [20XUAS-IVS-GCaMP6f}attP40/+; P{yw [GMR57C10-GAL4}attP2/+ (n=5)
2. w[+]; +; P{yw [GMR57C10-GAL4}attP2, PBac{w, 20xUAS-jRGECO1a-p10}VK5 (n=5)
3. w[+] norpA[36]; UAS-GCaMP6f / P{w[+mC]=ninaE-norpA.W}2; 39171-Gal4/+ (n=4)
4. w[−] norpA[36]; UAS-GCaMP6f / P{w[+mC]=ninaE-norpA.W}2; 39171-Gal4/+ (n=5)

Fly stocks were made using standard fly crossing techniques using balancer chromosomes and in the case of stock 2, recombination was used to bring two insertions onto the same chromosome. Note that P{yw [20XUAS-IVS-GCaMP6f}attP40 is abbreviated as UAS-GCaMP6f, P{yw [GMR57C10-GAL4}attP2 is abbreviated as 39171-Gal4. Stock 1 is referred to throughout as red eye/wild type photoreceptor function/GCaMP. Stock 2 is referred to throughout as red eye/wild type photoreceptor function/RGECO. Stock 3 is referred to throughout as red eye/Rh1 photoreceptors only/GCaMP. Stock 4 is referred to throughout as orange eye/Rh1 photoreceptors only/GCaMP. Females were used for stocks 1 and 2 and males for stocks 3 and 4.

### Fly Care

All flies were reared at 25-27°C with approximately 60% humidity under a 12:12 light/dark cycle and fed a standard cornmeal and molasses diet. Flies were collected one day after eclosion using CO2 for sedation and were transferred into fresh food vials. A prior study ^[42]^ cautions against using flies aged beyond 4-5 days if they carry the w1118 mutation for reasons of retinal degeneration, recordings were carried out in 4 to 7-day old flies. Slightly older flies were used due to ease of making the imaging preparation and did not show different results across this age range.

### Fly preparation

Following cold-anaesthesia (~15 min in a vial placed on ice), the fly was positioned on a sheet of aluminium foil such that the back of the head capsule protruded through a small cut-out and was tilted to form an approximate 10° angle with the foil sheet. This configuration allowed the back of the head to be exposed for imaging, while leaving the majority of the compound eye below the foil for visual stimulation. Using UV-curing adhesive, the fly was then secured to the sheet with special care to reduce brain movement by attaching the proboscis and legs to the upper thorax and immobilising the abdomen. A saline solution (103 mM NaCl, 3 mM KCl, 20 mM BES, 10 mM trehalose, 20 mM sodium bicarbonate, 1 mM sodium phosphate monobasic, 2 mM CaCl2 and 4 mM MgCl2, balanced to pH 7.4) was added to create a bath over the back surface of the head. The cuticle and underlying trachea were then gently removed to expose the optic neuropil. The trachea extending from the rim of the eye outward over the medulla was removed using forceps to gently pull it away. A gravity/suction pump was used to circulate saline over the course of the experiment to prevent build of up of metabolites or other resulting from the dissection. Following the removal of the cuticle, an interval of 45 minutes, or more, was allowed to ensure stabilisation of neural activity and dark adaptation of the fly.

### Visual stimuli

A projector (DepthQ 360 DLP, WXGA resolution) coupled to a screen was modified to use an independent light source, a monochromator (Cairn Research Optoscan Monochromator). This allowed us to project selectable narrow bands of the spectrum onto a screen material (Da-Lite, Polacoat^®^ Flex Plex Video Vision) that minimised wavelength-dependent scattering properties. Only the central portion of the screen was used, segmented by a circular ellipse of 400 pixels in diameter corresponding to 38.8 degrees of visual field. By removing the corners and edges, spectral variation resulting from light scattering is reduced (**Figure 2D**). The projector was used for patterned stimuli up to 360 Hz frame rate as some fly species, such as *Coenosia attenuata* can see up to 300 Hz, while *Drosophila melanogaster* can see just beyond 120 Hz ^[27]^. The light guide, the projector, the screen and the stage/holder were adjusted to ensure an ideal alignment of the optical pathway and visual field of the fly. A first adjustment was made to the light guide and projector mirror position to place the maximal brightness at the screen projection centre with an even drop in optical power towards the edges of the screen in all directions. Next, the holder was positioned in relation to the screen such that the holder centre is perpendicular to the projection centre and equidistant to all four corners. Furthermore, the holder and screen/projector are adjusted such that when the holder is positioned to be parallel to the screen, the spectrometer samples light directly at the centre of the screen. A motorised tip/tilt system allowed us to precisely tilt the holder in any direction such that the central point of the holder where the spectrometer detector (or fly’s eye) is positioned remains in the same point of space but the angle of the detector varies such that it samples light from a different location of the screen. Radiance spectra were measured using a NIST calibrated Avantes AvaSpec 2048 Single Channel spectrometer coupled to an Avantes UV-VIS 600 μm fibre (numerical aperture NA = 0.22, acceptance angle = AA = 25.4° and solid angle SA = 0.1521°).

#### Intensity-Response Relationship

The first set of visual stimuli applied to the fly were used to establish intensity-response relationships of medulla neurons. The stimulation protocol consisted of three repeated 1-second light flashes intercalated with 3 seconds of darkness. This motif was repeated for a range of twenty incremental intensities spanning four log units of light (from 10^8^ to 10^11^ photons/s/m^2^) to establish the dynamic range of the visual response. Intensity-response relationships were probed for three different bands of the spectrum defined by the following centre wavelengths: 385 nm (UV), 440 nm (blue) and 565 nm (green). These colours were chosen with the intent to match the known spectral sensitivity peaks of the fruit fly opsins within the limitations of the system, such as the lack of UV light transmission through the projector and the restriction of most green wavelengths by the filters to prevent sampling by the GaAsP detectors.

#### Spectral Response

Next, in order to probe spectral response properties, the fly was presented with a spectral sweep of randomised light flashes of varying centre wavelengths ranging from 390 to 720 nm, all calibrated to produce the same radiance of 2.63 × 10^9^ photons/s/m^2^. This intensity was chosen as it sits in the middle of the intensity-response range across all tested wavelengths and all tested fly genotypes. A gap in the spectral sweep was introduced spanning 490 to 656 nm and 595 to 700 for the green-emitting GCaMP and the red-emitting RGECO, respectively, to accommodate the red and green GaAsP detector sensitivities. Flashes of light were applied for 1 second with a 3-second interval between each pulse, and the entire spectral sweep was repeated five times with a different colour randomisation for each repetition. Spacing between centre wavelengths was originally set to be 5 nm increments, as specified by the voltages supplied by the monochromator software. However, an assessment of the Gaussian curve describing the band of light revealed that the peak of each curve is, in fact, not aligned with the expected centre wavelength, but shifted in either direction by a negligible amount. Nonetheless, in the interest of precision, centre wavelengths have been adjusted in graphs throughout.

#### Further visual stimulus considerations

Daily calibrations were carried out for all stimuli, and a post-calibration check was conducted to ensure precision of the stimuli. Each experiment consisted of all stimuli types and these were always presented in the following order: spectral sweep, green-, UV- and blue intensity-response stimuli. A ten-minute dark adaptation period preceded the five repetitions of the spectral sweep and preceded each presentation of the intensity-response stimulus.

### Two-photon imaging

Calcium signals in neurons of the medulla were imaged using a two-photon Bruker (Prairie Technologies) In Vivo Microscope using GFP and RFP detection channels, with a 20X water immersion objective (Zeiss, W Plan-Apochromat 20x/1.0 DIC M27; Cat # 421452-9600-000), modified with the addition of specialised optical bandpass filters (**Figure 1B** and **Supplementary Figure 1S**). An insight DS+ laser (Newport Spectra-Physics), with 920 nm infrared excitation applied to the sample. For functional imaging of calcium responses, data was acquired at a 512 x 512-pixel resolution at a rate of approximately 30 frames per second. Visual stimuli were generated in StimGL (Howard Hughes Medical Institute) and controlled via Matlab (Mathworks, MA, USA). The start of a visual stimulus sequence was indicated to the imaging software (PrairieView) by a trigger, which subsequently initiated the acquisition of images, but also controlled the monochromator and projector. Images were acquired using the green channel (GaAsP detector 1) for flies expressing GCaMP and the red channel (GaAsP detector 2) for flies expressing RGECO. A 200X zoom (20X on the objective and 10X on the software) was used to visualise the fly medulla. This allowed imaging of all layers of the neuropil from proximal to distal across a number of cartridges (ranging from a half a dozen to a dozen in different experiments, 0.104 to 0.115 μm/pixel with a field of view of 53 to 59 μm).

### Analysis

#### Fluorescent signal extraction

Analyses were conducted using custom-made scripts in Matlab (Mathworks, MA, USA). For region of interest (ROI) selection, a reference image was used corresponding to averaged frames across the response period for a blue light pulse. Circular ROIs 12 pixels in diameter were manually positioned to tile the sections of the 512 x 512-pixel image containing the medulla that are visually distinguishable from their fluorescence (both baseline and light response). The aim of this selection process was to assess responses in a given discernible layer without assuming response uniformity across the layer. Each ROI covers a sufficient number of pixels to ensure a good response could be extracted whilst limiting noise issues that may arise in larger ROIs. Each imaging sequence from a fly medulla averaged approximately 150 ROIs that were extracted for analysis. A cross-correlation analysis ^[47]^ was implemented to compensate for motion in the x-y plane resulting from physiologically driven movement of the visual neuropil (e.g. muscular displacement), vibrations resulting from equipment, or other. With this method, images across the experiment were realigned to an initial template image in order to ensure continuity of all features, and consequently continuity of ROIs from one image to another throughout the stack. A voltage recording paired to the two-photon image stack was used to segment the experiment into individual stimulus presentations. For each segment, baseline fluorescence was averaged over a pre-stimulus darkness period (5 frames prior to stimulus onset). Baseline fluorescence was averaged for each ROI over the pre-stimulus darkness period. Calcium activity was assessed by changes in fluorescence. These were calculated as the difference between average fluorescence for a given frame and the baseline fluorescence and then divided by the baseline for each ROI:

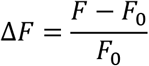

where *ΔF* is the change in fluorescence, *F* is the fluorescent signal and *F_0_* is the baseline fluorescence. Extracted responses were then averaged across stimulus repeats.

#### Curve fitting parameters and data selection

*ΔF* traces extracted for individual ROIs were averaged across repetitions of identical stimuli. In the case of data resulting from the spectral sweep, only three of the five repetitions were retained via a selection method whereby the two responses furthest from the mean of all five responses for a stimulus were removed. This methodology was introduced in an effort to reduce noise in the data. Next, a stimulus response value was calculated by integrating the response over time. The time segment for response integration was matched to the excitatory phase of the response profile observed to ensure maximal signal. In practical terms, integral values of the trace were calculated either over the one-second duration of the stimulus, over the one-second duration post stimulus or over the time course of both. Sigmoid curves were fitted to the intensity-response data using the Naka-Rushton equation classically used for electroretinogram data ^[48]^:

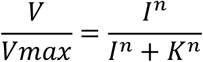

*V* is the response and *Vmax* the maximum amplitude of the response, *I* is the stimulus intensity, *K* is the intensity at the half-maximum response and *n* is the exponential slope measured at the half-maximum response. The curves were fitted in a two-step process. First, the intensity-response data was transformed for each stimulus intensity: 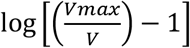. Then a linear regression was fitted to the transformed data to determine an initial approximation of *K*and *n* ^[49, 50]^. Next, the estimated *K* and *n* values were used to fit the curve with the above equation. The goodness-of-fit of the curve was evaluated by means of the mean square error value (MSE) and curves presenting MSE values superior to 0.05 were discarded. Furthermore, data was retained only if a curve could be fitted to all three intensity-response stimuli colours (UV, blue and green) within the set MSE parameter threshold.

A selection process was also applied to spectral sweep data. Spectral response curves were only retained for a given ROI if the following conditions were fulfilled: (1) all three intensity-response curves were fit as per the above criteria and (2) the light intensity used for the spectral sweep sits at least 1/5th below the maximal value of the intensity-response curve for all three colours (to ensure saturation was not reached at any point of the spectral sweep).

#### Statistics

All stages of data analysis (image analysis, curve fitting, data representation and statistical analyses) were conducted using custom scripts which used statistical packages available in Matlab (Mathworks, MA, USA). Individual statistical tests are reported in figure legends and the results of post hoc tests, original data points, mean, standard deviation, and standard error of the mean are reported throughout using notBoxPlot Matlab function (version 1.31.0.0, Rob Cambell).

## Acknowledgements

TJW held a Biotechnology and Biological Sciences Research Council (BBSRC) David Phillips Fellowship (BB/L024667/1) and RCF was on the BBSRC Doctoral Training Partnership. We thank the University of Cambridge for supporting the partial funding of our system and space to conduct our research. We also thank Bruker Fluorescence Microscopy (particularly John Rafter) for their support in integrating optics to their system, Semrock for assisting with custom fabrication of high quality optics, Cairn Research (particular Jeremy Graham) for assistance with customisation of the optoscan monochromator, HHMI Janelia farm for specifications to modify the DepthQ projector to accept a light guide source, the University of Cambridge fabrication workshops and Milly Sharkey for assistance with monochromator calibration. We also thank Mathias Wernet for anatomical insights and the many people who helped with other resources and comments both along the way and for final manuscripts.

## Author information

T.J.W. developed the initial study idea, designed and built optics system and generated transgenic *Drosophila* lines. R.C.F. conceptualised the monochromator light stimuli and projector screen system. T.J.W. and R.C.F. designed the experiments. R.C.F. conducted the experiments, analysed the data and wrote the manuscript. T.J.W. and R.C.F. finalised the manuscript.

## Additional information

### Competing interests

The authors declare no competing interests

## SUPPLEMENTARY INFORMATION

**Figure S1.**
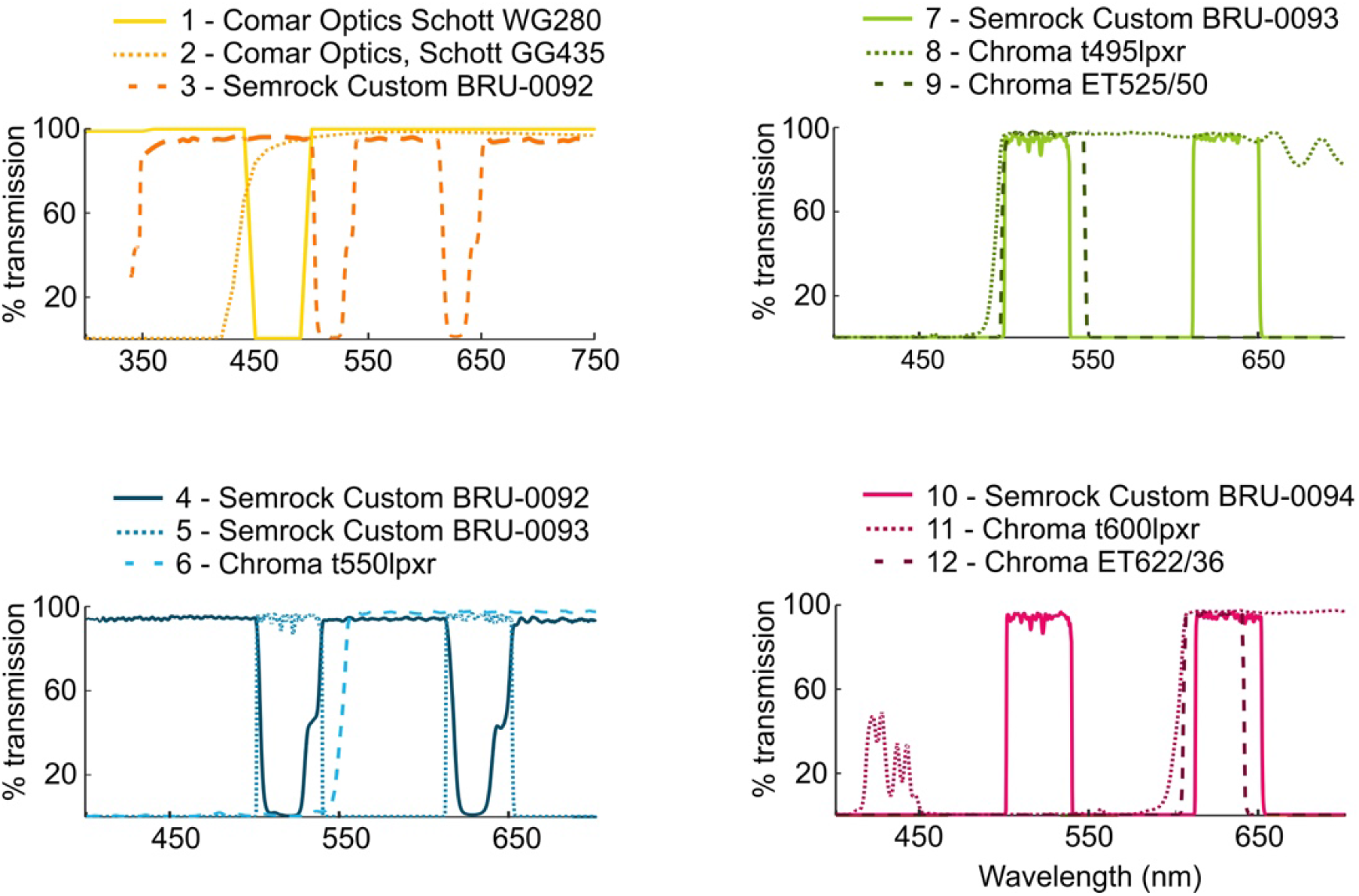
Filter spectra for the monochromator and the microscope. Filter spectra for the modified optical pathways in the monochromator and the microscope depicted in **Figure 1B**.

**Figure S2.**
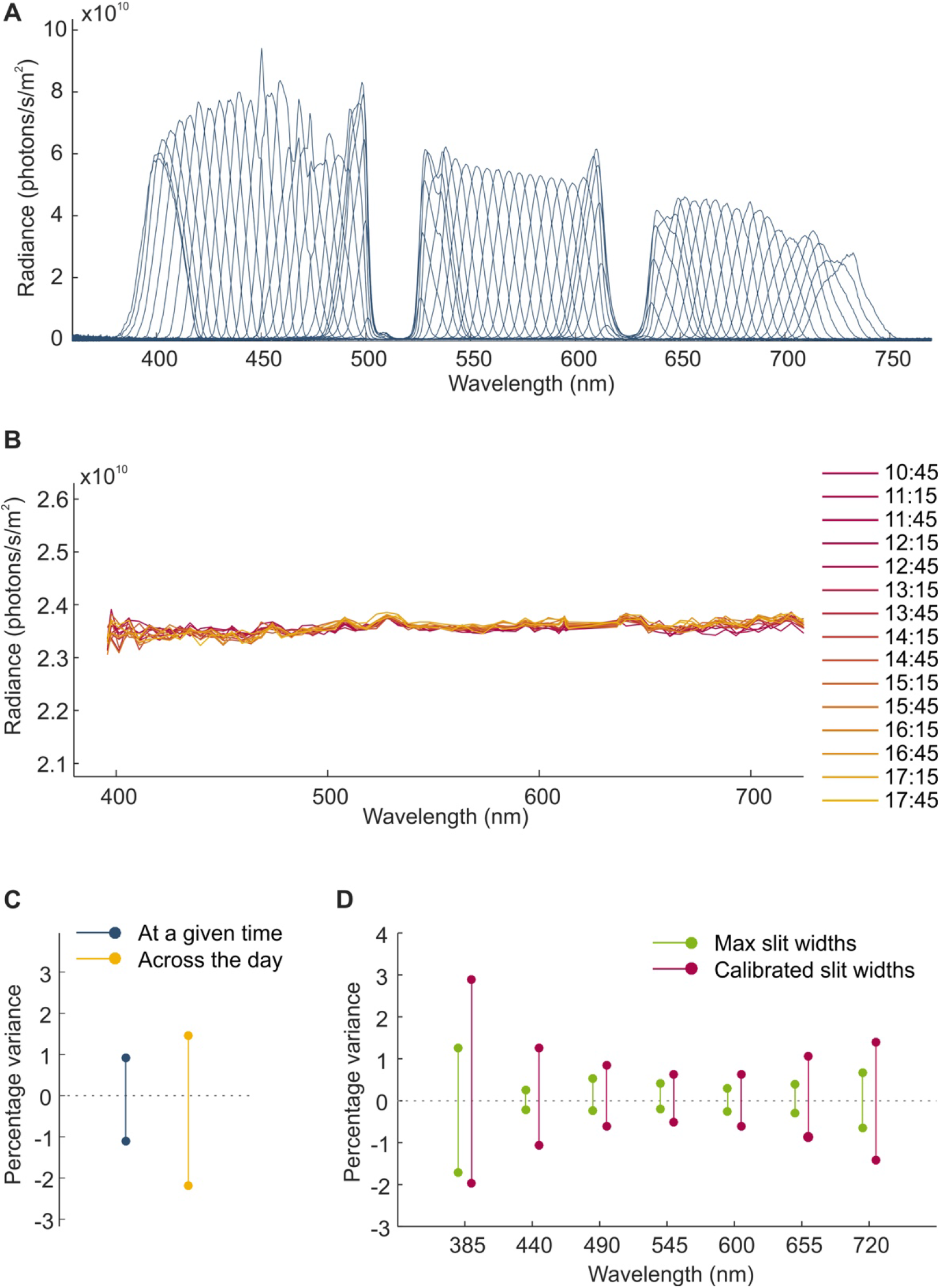
Detailed optical specifications of the monochromator-projector setup. (**A**) Radiance curves for each centre wavelength (385 to 720 nm in approximate 5 nm increments) for monochromator input and exit slits calibrated to produce equal brightness corresponding to a set reference value (here the radiance of the dimmest band of light, 385 nm centre wavelength, for a 30 nm slit width). (**B**) Calibrated radiance values determined from the area under the curve for each radiance spectra, measured at half hour intervals across the experimental day. (**C**) Brightness fluctuation of calibrated light across the spectrum reported as the maximum and minimum percentage variance from the mean at any given time (see B), and over the course of the entire day. (**D**) Power fluctuations for individual centre wavelengths at maximum slit width and calibrated slit widths over the course of 5 minutes. Percentage change corresponds to maximum and minimum change from the mean of measurements taken every second over the course of five minutes.

**Figure S3.**
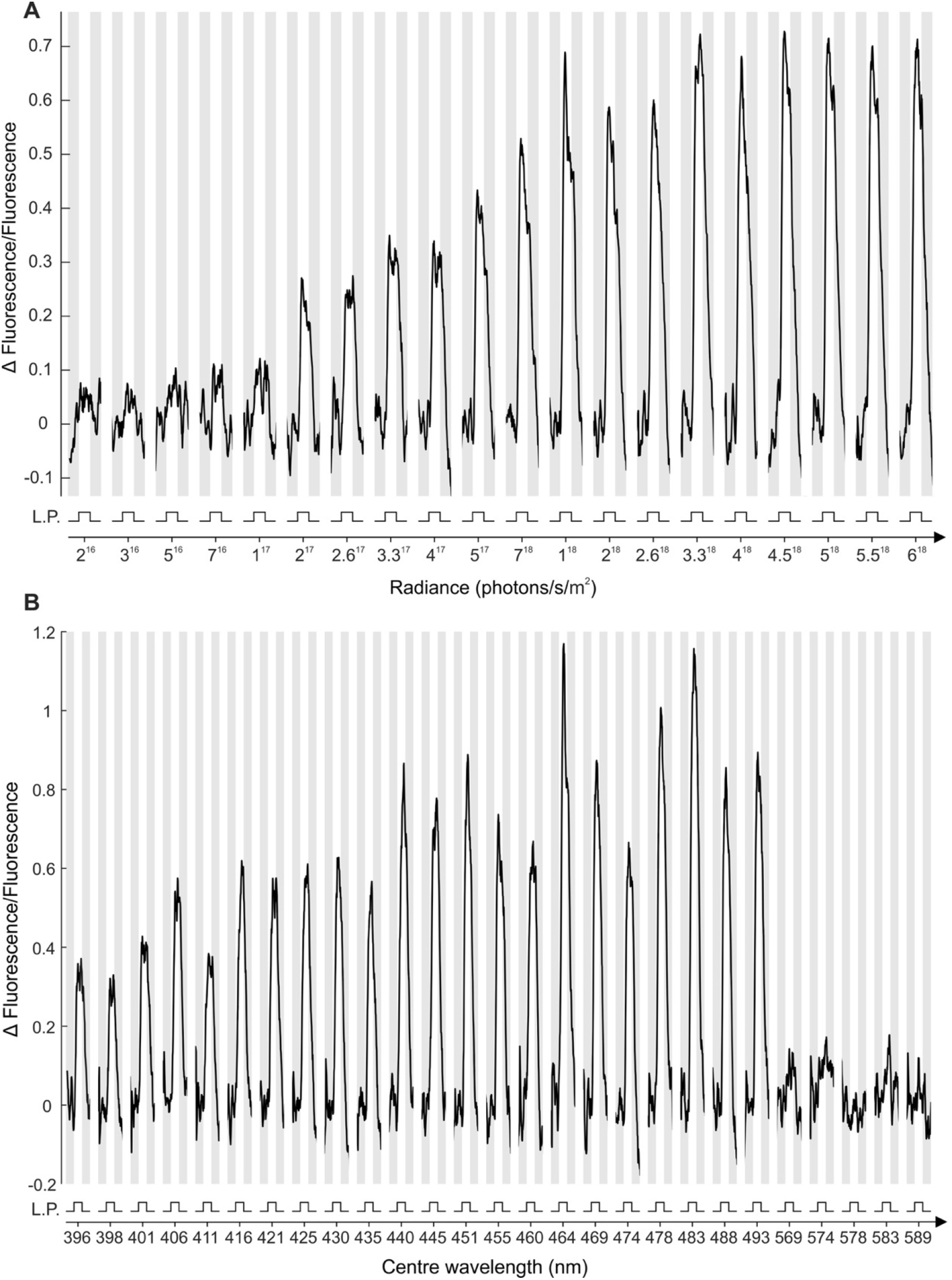
Intensity-response and spectral sweep stimuli: example responses. (**A**) Example fluorescence responses of an ROI to the intensity-response stimulus protocol consisting of light pulses (L.P.) of increasing intensity. Traces for a given intensity represent the mean of the response from all three stimulus repeats. (**B**) Example fluorescence responses of an ROI to the spectral sweep stimulus protocol consisting of light pulses of varying centre wavelengths. Traces for a given intensity represent the mean of the response from all three stimulus repeats. Response traces are not represented beyond 590 nm for the sake of clarity as no discernible change in fluorescence is detected beyond this point. Note that pulses were presented randomly but have been reordered here in ascending wavelength.

**Figure S4.**
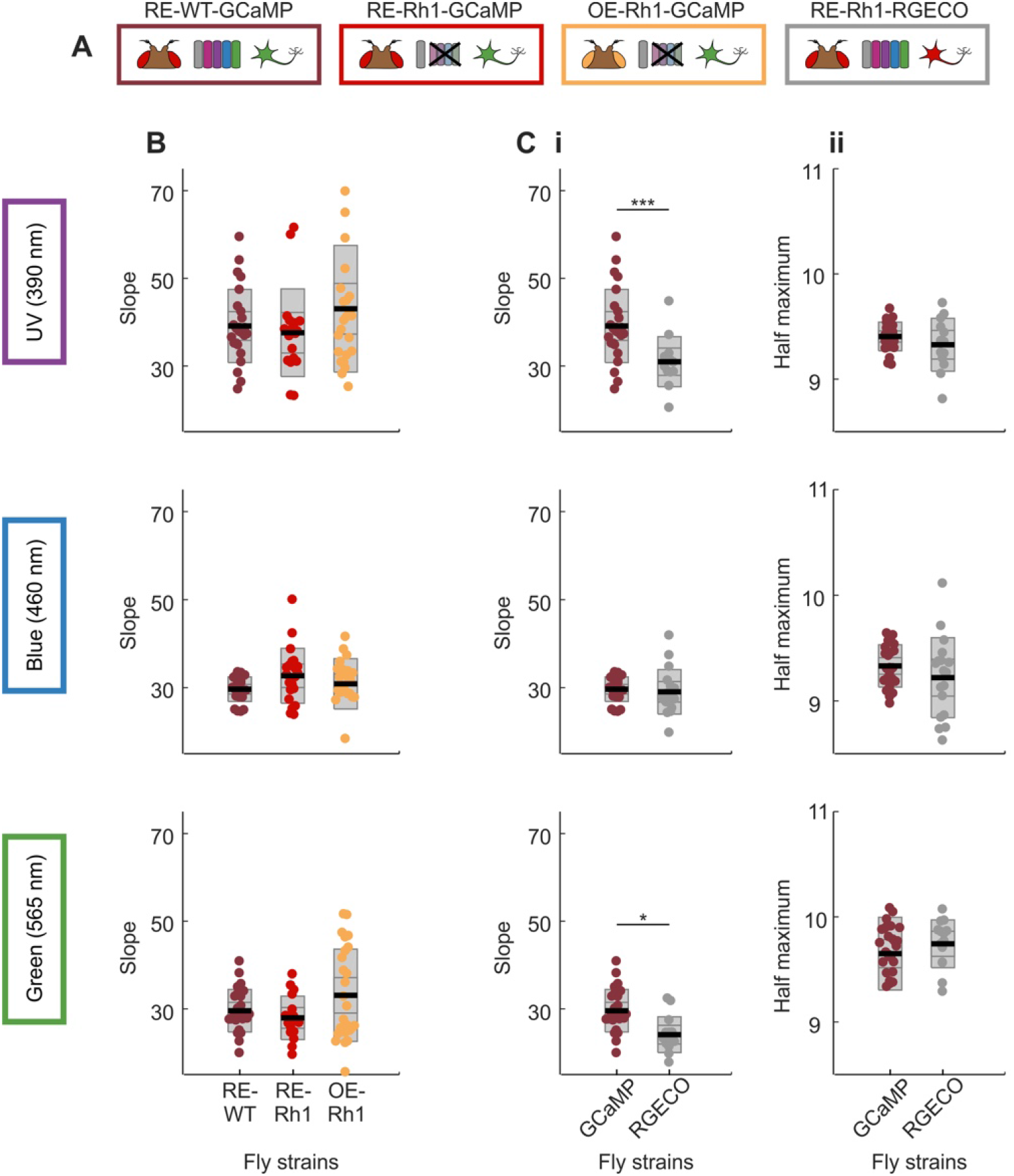
Intensity-response relationship coefficient comparison across fly strains. (**A**) Experiments were performed in four different fly strains expressing pan-neuronal calcium activity indicators– red eye, wild type photoreceptors and GCaMP6f (RE-WT-GCaMP, bordeaux); red eye, Rh1 only and GCaMP6f (RE-Rh1-GCaMP, red); orange eye, Rh1 only and GCaMP6f (OE-Rh1-GCaMP, orange) and red eye, wild type photoreceptors and RGECO (RE-WT-RGECO, grey). (**B**) Slope values extracted from intensity-response relationship curves for three different bands of monochromatic light (UV, blue and green; centre wavelengths 390, 460 and 565 nm respectively) for GCaMP-expressing fly strains. (**C**) Slope (**i**) and half maximum (**ii**) values extracted as in (B) comparing responses in red eye/wild type photoreceptors flies expressing either GCaMP6f or RGECO. Individual data points correspond to the average of ROI responses across a given layer structure (see methods) for a given fly preparation. The curves in (A) are a mean of the fitted curves of individual data points. Black line = mean, inner grey box = SEM, outer grey box = SD. Significant differences are noted with star values, with the P values from top to bottom: ***P = 0.0005 and *P = 0.0415 (one-way ANOVA).

**Figure S5.**
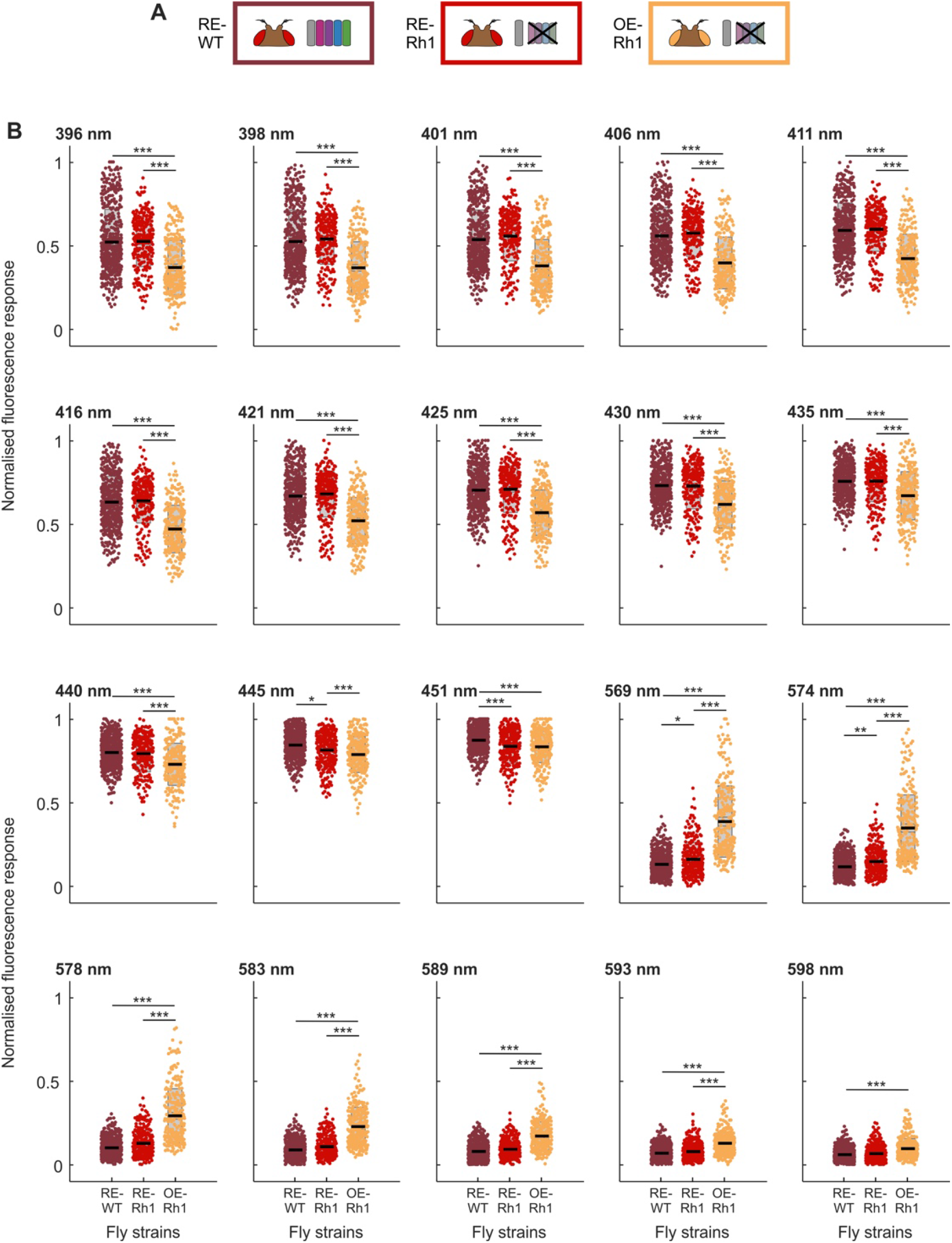
Spectral response profiling – breakdown for individual wavelengths. (**A**) Fly strains expressing pan-neuronal GCaMP6f– red eye/wild type photoreceptors (RE-WT, bordeaux); red eye/Rh1 only (RE-Rh1, red) and orange eye/Rh1 only (OE-Rh1, orange). (**B**) Responses to the individual wavelengths of the spectral sweep in Figure 3D that exhibit significant differences. Black line = mean, inner grey box = SEM, outer grey box = SD. Significant differences are noted with star values, with one-way ANOVA P values reported in table S1.

**Figure S6.**
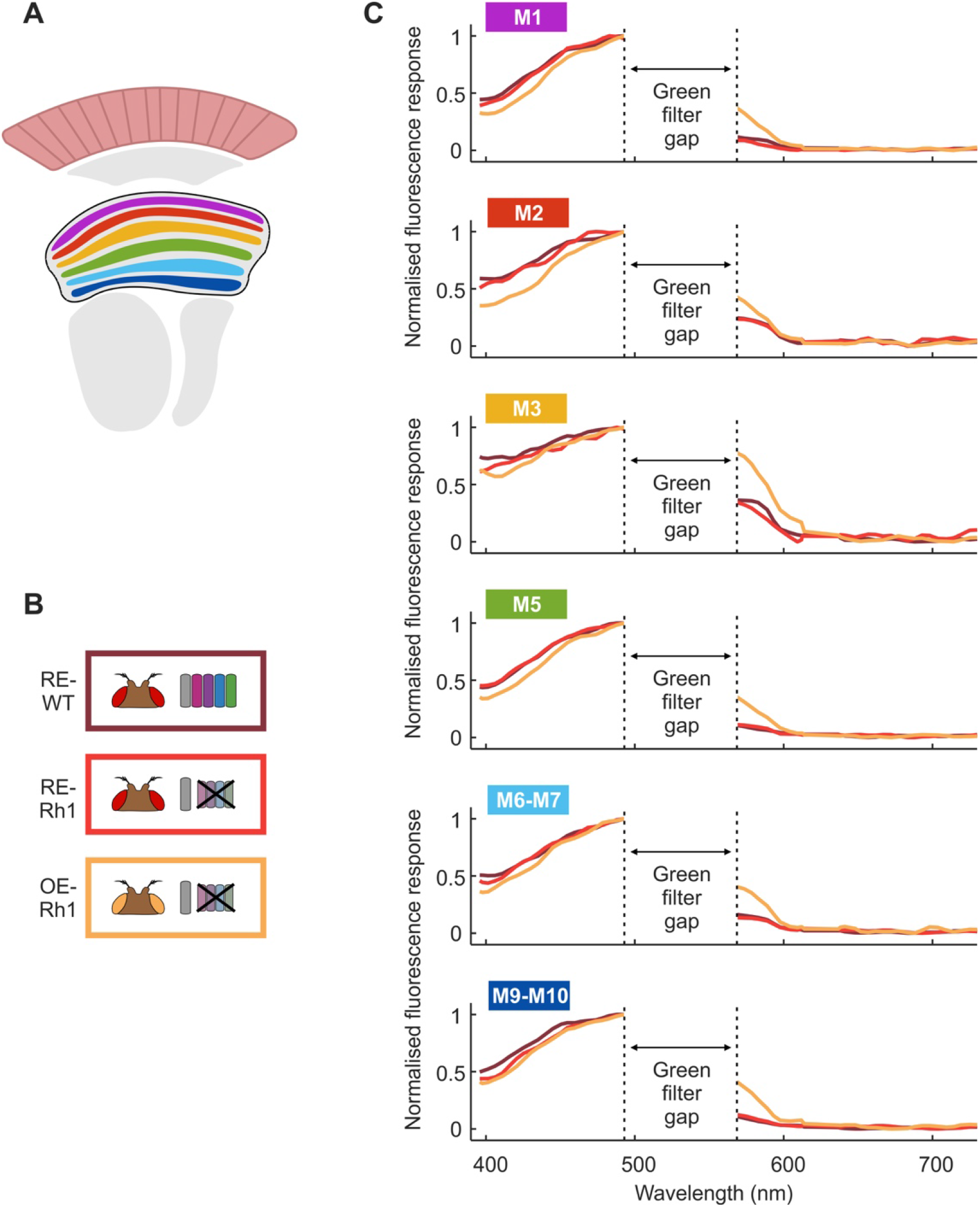
Spectral response profiles of layer groupings in the medulla. (**A**) Schematic of layer structures in the medulla as distinguished from pan-neuronal GCaMP labelling (see Figure 4). (**B**) Fly strains expressing pan-neuronal GCaMP6f– red eye/wild type photoreceptors (RE-WT, bordeaux); red eye/Rh1 only (RE-Rh1, red) and orange eye/Rh1 only (OE-Rh1, orange). (**C**) Mean spectral response profiles for layers M1, M2, M3, M5, M6-M7 and M9-M10.

**Table S1.** 1-way ANOVA p-values from Figure 3D and Figure S5B.

|  | RE-WT/RE-Rh1 | RE-WT/OE-Rh1 | RE-Rh1/OE-Rh1 |
| --- | --- | --- | --- |
| **396** | 1 | 3.2094e-05 | 3.2094e-05 |
| **398** | 1 | 3.2094e-05 | 3.2094e-05 |
| **401** | 0.8996 | 3.2094e-05 | 3.2094e-05 |
| **406** | 0.9999 | 3.2094e-05 | 3.2094e-05 |
| **411** | 1 | 3.2094e-05 | 3.2094e-05 |
| **416** | 1 | 3.2094e-05 | 3.2094e-05 |
| **421** | 1 | 3.2094e-05 | 3.2094e-05 |
| **425** | 1 | 3.2094e-05 | 3.2094e-05 |
| **430** | 1 | 3.2094e-05 | 3.2094e-05 |
| **435** | 1 | 3.2094e-05 | 3.2094e-05 |
| **440** | 1 | 3.2094e-05 | 3.2094e-05 |
| **445** | 0.0294 | 0.6676 | 3.2094e-05 |
| **451** | 1.9732e-04 | 4.3811e-05 | 1 |
| **569** | 0.0283 | 3.2094e-05 | 3.2094e-05 |
| **574** | 0.0098 | 3.2094e-05 | 3.2094e-05 |
| **578** | 0.1550 | 3.2094e-05 | 3.2094e-05 |
| **583** | 0.9891 | 3.2094e-05 | 3.2094e-05 |
| **589** | 1 | 3.2094e-05 | 3.2094e-05 |
| **593** | 1 | 3.2094e-05 | 3.2309e-05 |
| **598** | 1 | 2.5220e-04 | 0.2610 |

**Table S2.** 1-way ANOVA p-values from Figure 4C.

|  |  | RE-WT/RE-Rh1 | RE-WT/OE-Rh1 | RE-Rh1/OE-Rh1 |
| --- | --- | --- | --- | --- |
| **M1** | UV | 1 | 1 | 1 |
|  | Blue | 1 | 0.0039 | 0.6421 |
|  | Green | 1 | 7.2533e-06 | 7.2533e-06 |
| **M2** | UV | 7.2533e-06 | 1 | 7.2533e-06 |
|  | Blue | 7.2533e-06 | 0.7172 | 7.2533e-06 |
|  | Green | 7.2533e-06 | 7.2533e-06 | 7.2533e-06 |
| **M3** | UV | 0.9962 | 1 | 0.9960 |
|  | Blue | 0.4919 | 1 | 0.9999 |
|  | Green | 0.9957 | 0.3603 | 1 |
| **M5** | UV | 1 | 0.1753 | 0.9403 |
|  | Blue | 1 | 1 | 1 |
|  | Green | 1 | 7.2534e-06 | 7.2534e-06 |
| **M6-M7** | UV | 1 | 0.0335 | 0.9671 |
|  | Blue | 1 | 0.2098 | 0.9167 |
|  | Green | 1 | 1 | 1 |
| **M9-M10** | UV | 1 | 0.0022 | 0.8434 |
|  | Blue | 0.9999 | 1 | 0.9981 |
|  | Green | 1 | 7.2534e-06 | 7.2534e-06 |

## Notes

### Competing Interest Statement

The authors have declared no competing interest.

### Summary of Updates

Based on a HH journal editor feedback, that this was technique was not novel and lacked color responses from single neurons downstream of the receptors, we have refined the text to highlight that many aspects of our method are novel and what we have actually done has characterised the system and shown novel findings.

